# Food environment components influencing consumption trends of neglected and underutilized species in northern Benin

**DOI:** 10.1101/2024.11.13.623353

**Authors:** Bissola Malikath Bankole, Waliou Amoussa Hounkpatin, Sam Bodjrenou, Julia Bodecker, Lynn Julen, Flora Josiane Chadare, Celine Termote

**Affiliations:** School of Nutrition, Food Science and Technology, Faculty of Agronomics Sciences, University of Abomey-Calavi, Benin; Alliance Bioversity International – CIAT, Benin; Alliance Bioversity International – CIAT, Kenya; Laboratory of Science and Technology of Food and Bioresources and Human Nutrition, Sakété University Center, National University of Agriculture, Benin

**Keywords:** Food environment, neglected and underutilized species, food systems, drivers, Benin

## Abstract

Malnutrition is a global problem driven by food systems that impact climate and biodiversity. Neglected and underutilized species (NUS) could improve diets, but what drives their choices and consumption, particularly in low and middle-income countries, is poorly documented. This study investigated the influences of the personal food environment on the consumption of NUS in three communities in the department of Atacora in northern Benin. Following a purposive sampling strategy, 24 semi-structured interviews were conducted with local and 18 with professional experts. 12 group discussions with villagers from six villages in the three communes complemented and deepened the information gathered, focusing on both barriers and factors for enhancing consumption. The data collected was translated and transcribed into French and analyzed using qualitative content analysis with Atlas.ti software. Among the investigated plant parts, an upward trend was found for *Moringa oleifera* leaves and *Vigna radiata* seeds, a downward trend for *Adansonia digitata* pulps, and a varied trend for *Ocimum gratissimum* leaves and *Adansonia digitata* kernels and leaves. Drivers for changes in consumption were found in all four dimensions of the food environment. Among the plant parts with the main increasing trend in consumption, various aspects of desirability, above all increased food and nutrition knowledge and skills, led to a positive consumption trend. The downward trend has most often been attributed to declining accessibility, but several aspects have also made these plant parts less affordable and less desirable (taboos for example). The strong variation in dimensions’ influence on plant parts with variable trends reflects their non-unanimous changes. Research and policies should address the factors influencing the consumption of these foods. Making neglected species more accessible, affordable, and desirable can enhance food security and environmental sustainability.

## 1. Introduction

A sustainable food system is defined as one that guarantees food security and nutrition for all so that the economic, social, and environmental bases for generating food security and nutrition for future generations are not compromised (1). Greater use of agrobiodiversity in food systems is a promising approach to help achieve complex, interconnected goals that span nutrition, environment, and health (2). Today’s food systems, dominated by few, highly uniform crops, are detrimental to our nutrition, health, and environment, as well as to the cultural identity of populations, their autonomy, and the economic empowerment of communities engaged in agriculture and vulnerable groups, including women (3). In terms of healthy and nutritious food needs, the potential of many edible plant species remains untapped (4,5). Neglected and underutilized species (NUS) encompasses wild, semi-domesticated, or fully domesticated plants in various forms such as field crops, trees, shrubs, and vines, along with edible fungi and animal species whose potential is not fully exploited (6). Many NUS have similar or better nutritional profiles than most common foods (3). NUS can enhance the accessibility of healthy foods in rural regions through various means: enhancing the availability of nutritious foods for direct consumption by farmers; reducing the expenses associated with obtaining a healthy diet, particularly wild foods; boosting the presence of nutritious foods in local food stores and markets for sale by producers; and raising income levels by tapping into niche marketing opportunities, which can subsequently be utilized to buy other, more nutritious foods (7). Concerning food and nutritional security, locally available food resources, whether cultivated or wild, can contribute to human nutrition by providing sources of diet diversification. They are rich in energy, micronutrients, vitamins, and minerals, and could offer solutions in combating ‘hidden hunger’(8,9). NUS can also be valuable sources of protein, fiber, and bioactive compounds, such as flavonoids, that significantly contribute to dietary health (10).

Despite their potential, various geographical, agronomic, economic, and socio-cultural aspects in food system value chains represent barriers to NUS consumption (6,11). More specifically, these barriers are mainly attributed to the poor economic competitiveness of NUS compared to staple crops, inefficiencies in the production, storage, and processing of NUS, the lack of sound baseline data on the nutritional and protective properties of NUS, disorganized or nonexistent food supply chains, negative associations with a poor rural lifestyle and low social status (11). Thus, food choice and consumption are influenced by various physical, economic, social, and cultural factors (12) and are dynamic, contextual, multifaceted, and multilevel (13).

Food environments serve as the interface for individuals to interact with the larger food system for food acquisition and consumption (14). The internal domain consists of individual-level dimensions of accessibility, affordability, convenience, and desirability whereas the external domain includes availability, prices, vendor and product properties, and marketing and regulation (15). In the rural food environments of low-income countries, food consumers are often food producers, and many foods are produced and obtained as part of the local food system (7). Their food environment is inherently connected to the local production system and the well-being of the local ecosystem. Thus, food is often acquired through the natural and built food environment, including market acquisition, crop harvesting, and wild collection, as well as through non-monetary transfers (16,17). Reintroducing NUS into food environments could improve access to and utilization of nutritious foods, leading to healthier diets (3). There is a growing need to improve the use of neglected and underutilized species (NUS), which are undervalued by current food environments, even though they are often highly nutritious and resilient to climate change, and could offer new income-generating opportunities (9). A better understanding of local food environments in NUS, including drivers of consumption, is therefore essential to inform subsequent nutrition interventions to improve food and nutrition security, environmental security, and strengthening livelihoods (18). Thus, this study aimed to identify the personal food environment factors influencing the consumption of four targeted plant species (*Adansonia digitata, Moringa oleifera, Ocimum gratissimum and Vigna radiata*) by people living in three (03) of the seven (07) communes in the department with high levels of food insecurity: Natitingou with 27.8%, Tanguiéta with 26.5% and Toucoutouna with 29.8% of household’s global food insecure (19).

## 2. Materials and Methods

### 2.1. Sampling

In each commune, two villages were randomly selected namely Kampouya and Koukouarbirgou in the commune of Natitingou, Tectibayaou and Wabou in the commune of Toucountouna and Douani and Kosso in the commune of Tanguiéta. Purposive sampling (20) was used in this study to select informative interview partners in the area of interest. The actors to be included in each key informant category were identified by recommendation. Saturation theory was used to determine the sample size. Two groups of key informants were identified based on the following required information on the personal food environment and its influences on the consumption of the selected NUS.

- The first group was made up of local experts who were chosen based on the criteria of being at least 50 years old and with a longstanding history of living in one of the six selected research villages. The choice was made to ensure that all the ethnic groups in the area would be represented.
- The second group of key informants was made up of professional experts who were selected based on the criteria of their professional experience in agriculture, nutrition, and health with direct contact with the local population living within the three selected communes. These included members of advisory services, workers at social promotion centers, and local nutritionists working in the private or public sector.

Focus groups were also held with people from different ages, religious, educational and sociolinguistic groups, to validate and complement the information obtained from the semi-structured interviews. Participants in semi-structured interviews with local experts and village chiefs were excluded. Preferably, no participants from the same household were selected. To ensure the homogeneity of the focus groups, discussions with men and women were conducted separately to enhance the free exchange of opinions. Then, respectively 24 local experts and 18 professional experts were selected for this study while 12 focus groups were done. However, information was collected until saturation was reached.

### 2.2. Data collection

A pre-test was carried out to improve the different questionnaires. The first phase consisted of semi-structured interviews with local and professional experts from different perspectives on the personal food environment. Local experts were questioned about their personally perceived changes in the consumption of the selected NUS and related changes in their food environment while professional experts were asked about the changes they perceived while working with local community members within the research area. Then, for each NUS selected, an in-depth discussion was held on questions relating to all dimensions of the personal food environment.

Focus group discussions (FGD) took place in the second step. Questions were asked about the dimensions and aspects of food environments related to the four selected NUS. Examples of positive and negative influences of the factors drawn from the semi-structured interviews were integrated into the focus group questionnaire to deepen their understanding. During data collection, the same visual stimuli were used in the form of photographs of the selected NUS to make them easily identifiable. Photographs were numbered from 1 to 4 to facilitate handling of the different species (particularly for ranking tasks).

### 2.3. Data management and analysis

A total of 24 semi-structured interviews with local experts and 12 focus group discussions were recorded in local languages. All the recordings have been fully translated and transcribed into French.

#### 2.3.1. Ranking

Data saved were analyzed from a content perspective. Quantitative data from the rankings that originate from ordering photographs of the four NUS according to specific criteria were analyzed by descriptive statistics (frequencies). The scores cited by each respondent were added for each plant, resulting in a total score for each NUS. In the second step, this total is divided by the number of interview partners which resulted in a mean for each NUS. The lower the mean value, the higher was the rank of the species for the specific criterion (21). The overall mean rankings for the four species were then calculated from the ranked sums of the different interview groups (local and professional experts as well as focus groups).

#### 2.3.2. Qualitative content analysis

The first step of the qualitative content analysis involved coding, where relevant information from the 54 transcription documents was extracted according to the developed categories. The second step of content structuring consisted of paraphrasing, generalizing, and reducing the coded text passages and finally led to derived answers to the research questions. All text material was coded after the same procedure using the software ATLAS.ti version 7.5.18 (Scientific Software Development GmbH, Berlin, Germany).

#### 2.3.3. Coding

The unit of analysis consisted of 42 semi-structured interviews and 12 focus group discussions. Then, for each of the four (04) NUS, 13 distinct categories were created (Table 1). Categories 1 to 8 included relevant dimensions and aspects derived from the conceptual framework. Category 9, called “additional factors”, included relevant information influencing the consumption of the corresponding species, which could not be classified in one of the other existing categories. In addition, categories 10 to 13 described positive, negative, and neutral changes in the consumption of each species, as well as changes in consumption patterns over time and the present.

**Table 1:**
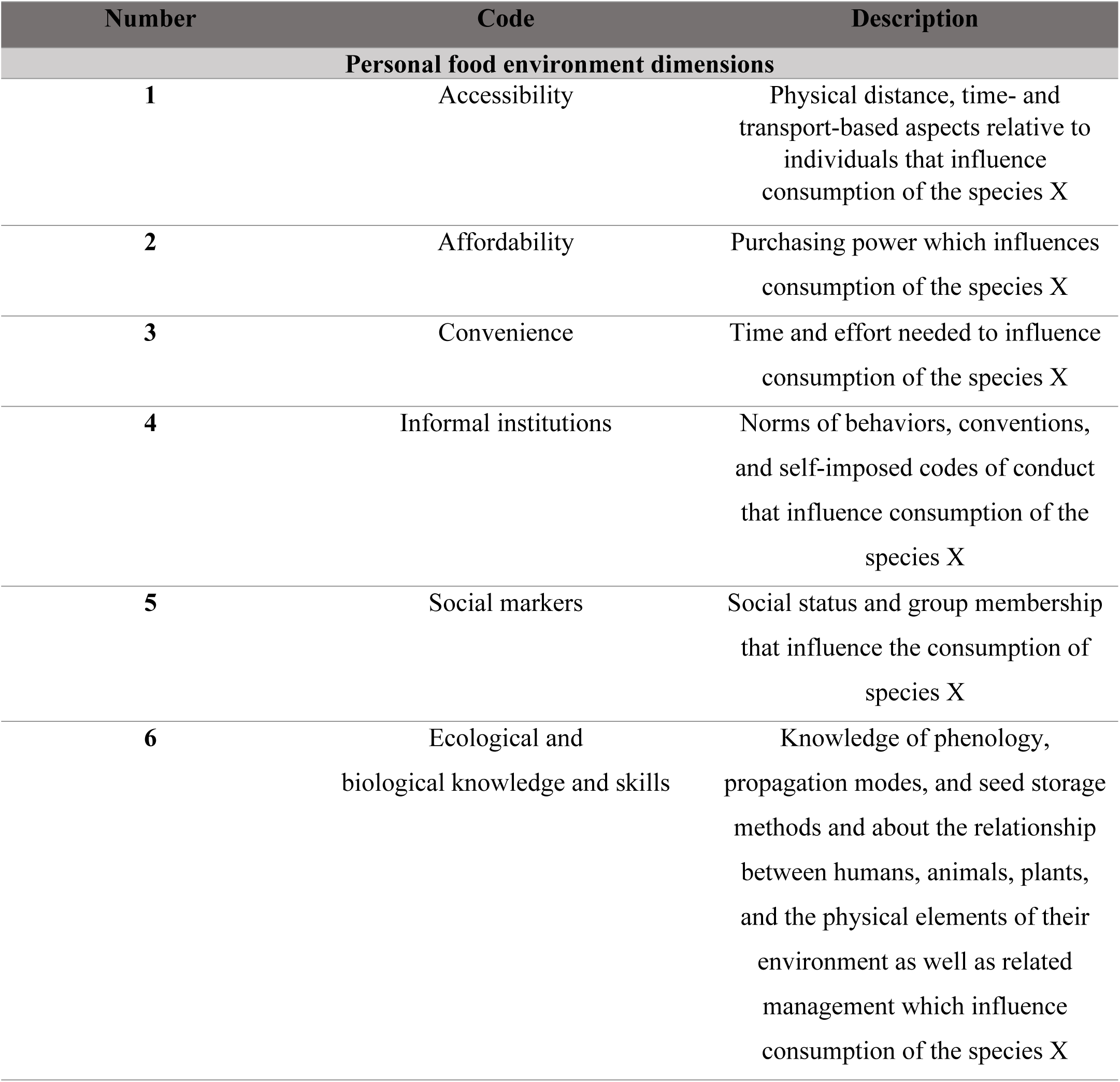
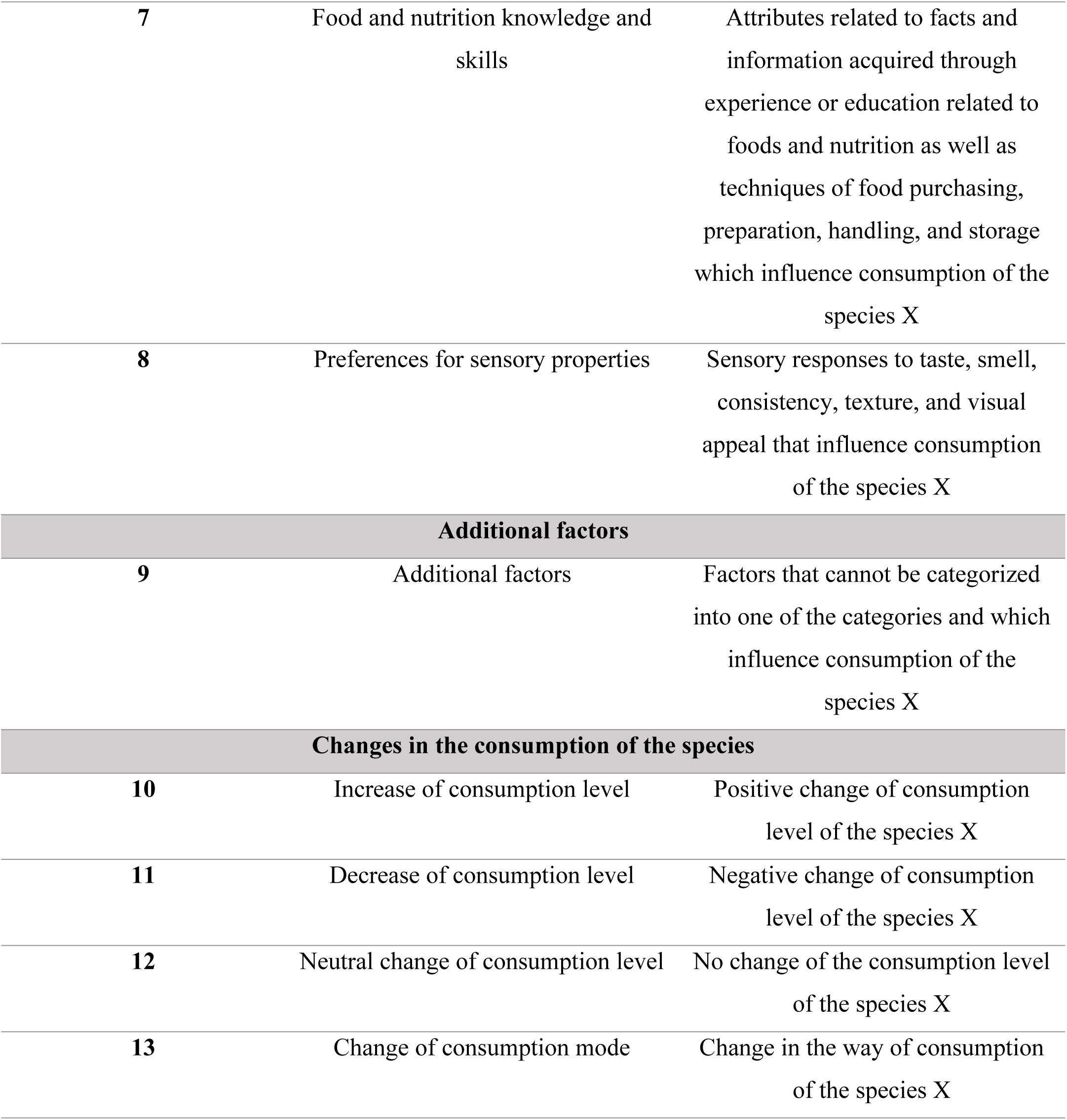
Category developed.

| Number | Code | Description |
| --- | --- | --- |
| **Personal food environment dimensions** |  |  |
| 1 | Accessibility | Physical distance, time- and transport-based aspects relative to individuals that influence consumption of the species X |
| 2 | Affordability | Purchasing power which influences consumption of the species X |
| 3 | Convenience | Time and effort needed to influence consumption of the species X |
| 4 | Informal institutions | Norms of behaviors, conventions, and self-imposed codes of conduct that influence consumption of the species X |
| 5 | Social markers | Social status and group membership that influence the consumption of species X |
| 6 | Ecological and biological knowledge and skills | Knowledge of phenology, propagation modes, and seed storage methods and about the relationship between humans, animals, plants, and the physical elements of their environment as well as related management which influence consumption of the species X |
| 7 | Food and nutrition knowledge and skills | Attributes related to facts and information acquired through experience or education related to foods and nutrition as well as techniques of food purchasing, preparation, handling, and storage which influence consumption of the species X |
| 8 | Preferences for sensory properties | Sensory responses to taste, smell, consistency, texture, and visual appeal that influence consumption of the species X |
| **Additional factors** |  |  |
| 9 | Additional factors | Factors that cannot be categorized into one of the categories and which influence consumption of the species X |
| **Changes in the consumption of the species** |  |  |
| 10 | Increase of consumption level | Positive change of consumption level of the species X |
| 11 | Decrease of consumption level | Negative change of consumption level of the species X |
| 12 | Neutral change of consumption level | No change of the consumption level of the species X |
| 13 | Change of consumption mode | Change in the way of consumption of the species X |

In the first step, the material was systematically filtered for information based on the category system. Information about characteristics regarding a specific code was extracted (increase/decrease/neutral change/changed). Some of the additional factors were added to one of the existing categories as they were identified as underlying drivers. The remaining additional factors included mainly statements related to physiological needs or general preferences/dislikes.

#### 2.3.4. In-depth analysis and content structuring

After paraphrasing and generalizing all statements, reduction encompasses the review of all generalized statements and was divided into two sub-steps. The statements with redundant information were reduced whereas the whole variation in information was kept. Firstly, the matrices resulting from the first reduction step included information on consumption trends and past and current consumption levels for each plant species or part studied. For that, the information from codes 10-13 (changes in the consumption of the species) was used and was quantitatively analyzed. Then, the respective statements were counted separately for semi-structured interviews and focus group discussions. The statements about increased/decreased/unchanged consumption were counted per focus group (unlike in semi-structured interviews where they were counted per person). Repeating statements by the same interview partner/focus group were counted once. A net change was calculated for each group by subtracting the number of declarations of decreased consumption from the number of declarations of increased consumption and dividing by the total number of declarations. The overall net change for consumption of each investigated plant part resulted from the difference of the total number of increased minus decreased statements divided by the total number of statements :

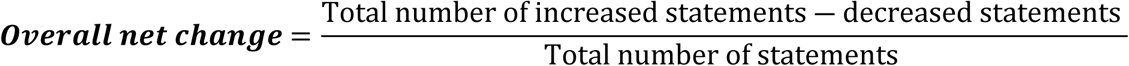

These numbers varying from +1 to -1 were classified into three main trends:

- increasing trend in consumption (1 to 0.33)
- varying trends in consumption (0.33 to - 0.33)
- negative trend in consumption (-0.33 to -1)

Then, information from the matrices of codes 1 to 9 was used to explain how the dimensions of the food environment can explain the different trends obtained for the different parts of the species consumed.

### 2.4. Ethics approval and consent to participate

This study was conducted according to the guidelines laid down in the Declaration of Helsinki. All procedures involving research study participants were approved by the Benin National Health Research Ethics Committee (CNERS-www.ethique-sante.org) with ethical approval number No.46 of November 07^th,^ 2019 and reference number No_093/MS/DC/SGM/DRFMT/CNERS/SA. Before each interview or group discussion, the project’s background and interview procedure were explained.

## 3. Results

### 3.1. Participant characteristics

All the local experts were farmers (100%) and the average age was 59 years (Table 2). The average duration they had lived in the research villages was 57 years. The interviewees belonged to the Waama (62.5%), Natimba (29.1%), and Otamari (8.3%) ethnic groups. Around half of those interviewed (54.1%) had received no formal education, while the others had attended elementary school. As the professional experts, more than half were men (66.6%) aged between 24 and 52 years. Most of them (61.1%) worked as nutritional advisors, followed by agricultural advisors (22.2%) and the others were social workers and scientists. The professional experts belonged to diverse ethnic groups. About 66.6% of the professional experts had a bachelor’s degree. Regarding the focus group, the respondents’ mean age was 41 years, ranging from 18-80 years. The mean residential duration in the research villages was 35 years, ranging between 6-72 years. All respondents were farmers. Religious affiliation and education level varied among participants but most of them were from Waama ethnic group.

**Table 2:**
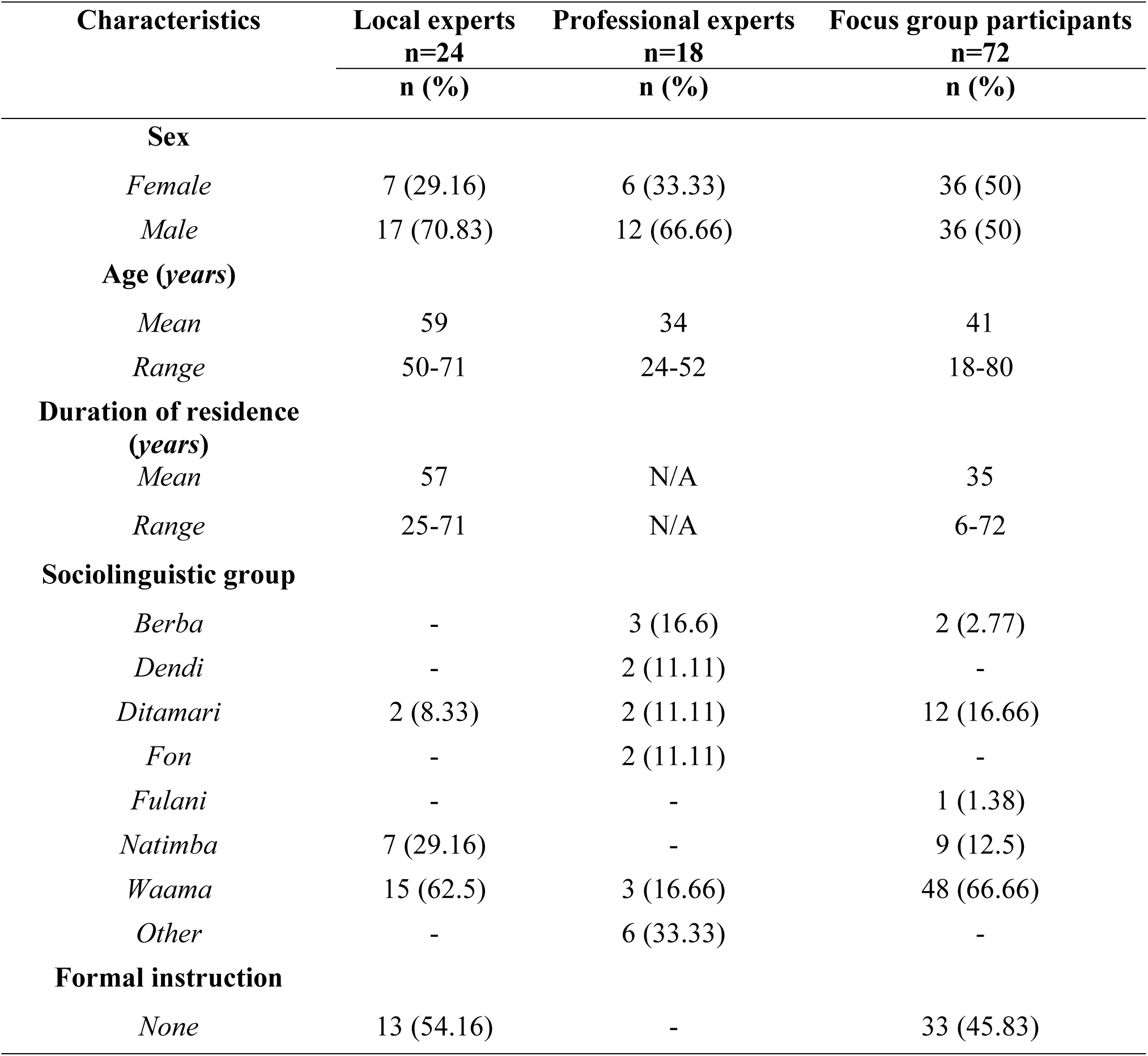
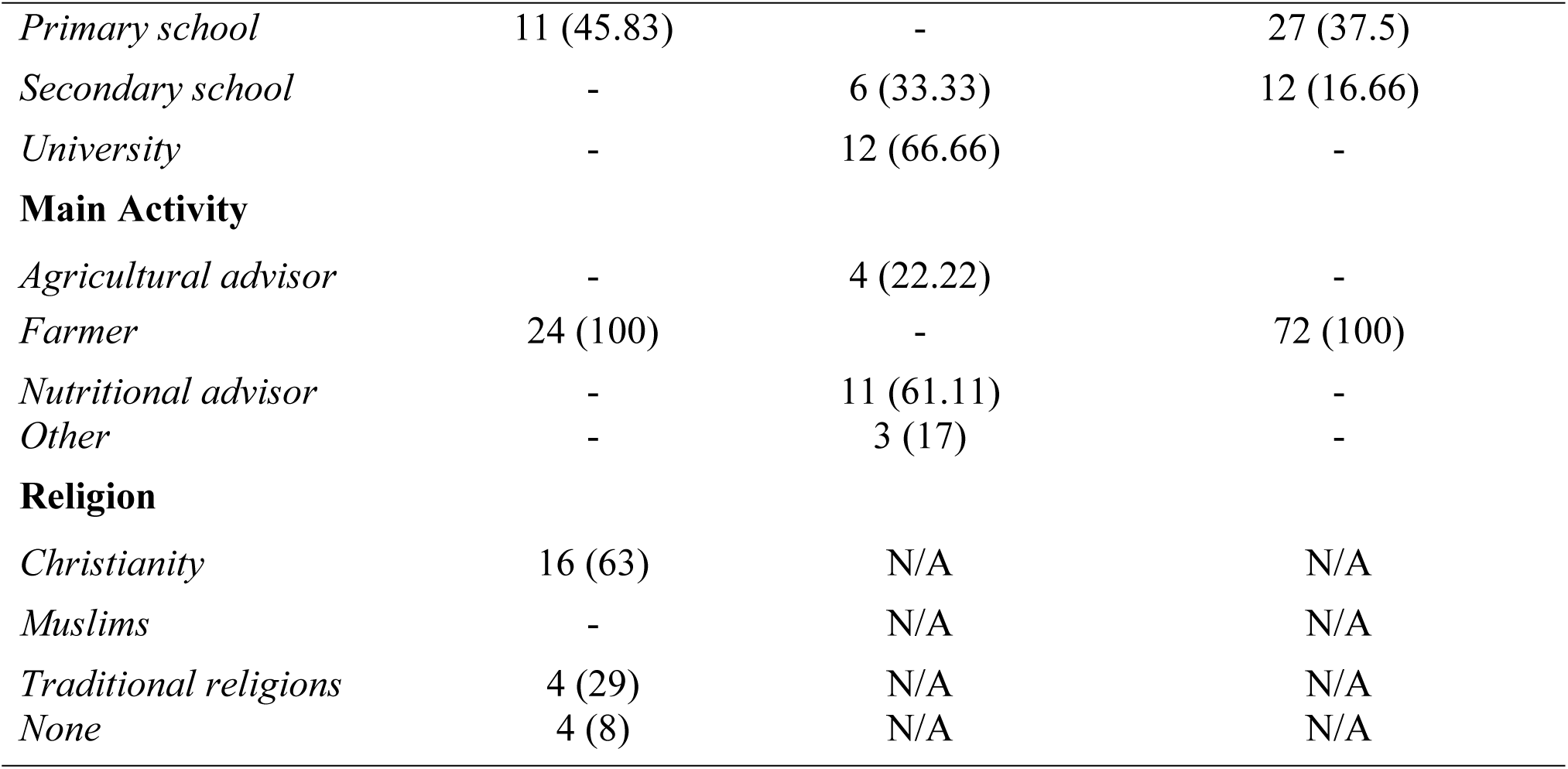
Sociodemographic characteristics of participants.

### 3.2. Consumption of NUS selected

Deeper insights in the relative consumption frequencies and preferences among the four (04) NUS were indicated through their rankings (Table 3). Most and least frequently consumed species were identical with the most and least preferred ones. *A. digitata* and *M. oleifera* were the most frequently consumed and preferred species, whereas *V. radiata* and *O. gratissimum* were on the lowest ranks.

**Table 3:**
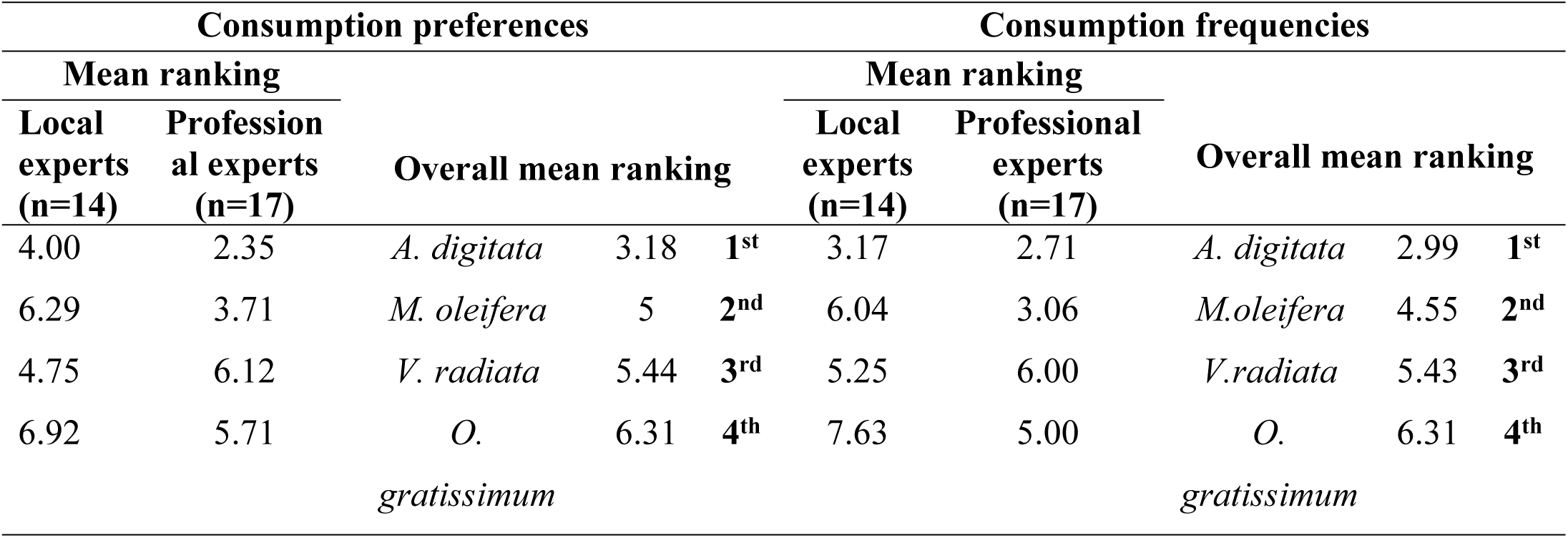
Mean ranking for consumption preferences and frequencies.

### 3.3. Changes in consumption of selected NUS

Changes in consumption over the last decades varied among the selected NUS as well as among different edible plant parts of the same species (Table 4). A major increase in consumption was predominant for *M. oleifera*’s leaves and *V. radiata*’s seeds. For *A. digitata* pulps, a downward trend in consumption was predominant. For the remaining investigated plant parts of four species (*A. digitata*’s leaves and kernels and *O. gratissimum*’s leaves), statements about their consumption varied in a more balanced way between increased, decreased, and unchanged consumption.

**Table 4:** Changes in consumption of the NUS plant parts.

| Species | Plant parts consumed | Number of statements by local and professional experts (N=42) |  |  |  | Number of statements by focus groups (N=12) |  |  |  | N=54 |  |
| --- | --- | --- | --- | --- | --- | --- | --- | --- | --- | --- | --- |
|  |  |  |  |  |  |  |  |  |  | Overall net | Main trend |
|  |  | Changes in consumption |  |  |  |  |  |  |  |  |  |
|  |  | Increase | Decrease | No change | Net change | Increase | Decrease | No change | Net change |  |  |
| *M. Oleifera* | Leaves | 20 | 1 | 1 | 0.86 | 5 | 0 | 0 | 1.00 | 0.89 | Increase |
| *V. radiata* | Seeds | 12 | 7 | 0 | 0.26 | 5 | 0 | 0 | 1.00 | 0.42 | Increase |
| *O. gratissimum* | Leaves | 4 | 5 | 2 | - 0.09 | 2 | 0 | 0 | 1.00 | 0.08 | Varied |
| *A. digitata* | Leaves | 4 | 3 | 3 | 0.10 | 2 | 0 | 1 | 0.67 | 0.23 | Varied |
| *A. digitata* | Kernels | 1 | 5 | 2 | -0.50 | 2 | 0 | 1 | 0.67 | -0.18 | Varied |
| *A. digitata* | Pulp | 3 | 11 | 3 | -0.47 | 1 | 0 | 1 | 0.50 | -0.37 | Decrease |

The clearest trend of increasing consumption was observed for M. oleifera leaves, whose consumption was non-existent or low a few decades ago. V. radiata seeds also showed a positive trend in consumption mainly linked to the recent period. They were traditionally known and consumed in the research area, although not all those interviewed said they had eaten the species in their youth. About ten years ago, consumption of V. radiata seeds fell to a virtually non-existent level. In recent years, there has been an increase in harvests from own production and higher levels of consumption, although some respondents still mentioned a total absence of seed consumption of the species currently. New methods of preparation include incorporating the seeds into traditional meals such as the Beninese dish “Atassi/Watché” (rice with beans), or bakery products such as cakes.

Among plant parts with a downward trend, consumption of A. digitata pulp has declined. A few decades ago, pulps were considered a sweetener and were often consumed directly by children or added to porridges.

For A. digitata leaves, positive, negative, and unchanged consumption trends were mentioned, with the level of commercialization of leaves having increased over recent decades. Others stated that consumption had only slightly increased, or even decreased since the leaves were already widely consumed in the past. The consumption of A. digitata kernels has undergone similar variations. The processing of A. digitata kernels into oil represented a new consumption pattern, in addition to their use in sauces. As for O. gratissimum leaves, their consumption was non-existent or low during the youth of those surveyed. Current consumption was limited to a minority of the local population but with varying trends over time.

### 3.4. Drivers for changes in consumption of the selected NUS

Multidimensional drivers within the personal food environment were identified as factors that contributed to the consumption trend of NUS.

#### 3.4.1. Moringa Oleifera

Regarding the **accessibility dimension**, although, some of those interviewed perceived no change in the accessibility of *M. oleifera,* as trees were already present nearby, growing in fields and around houses two decades ago, some changes have been noted: Indeed, improved accessibility has been achieved by increasing or emerging clean cultivation practices. *M. oleifera*, for example, started to be grown in home gardens. ‘‘*This species is produced around habitats and therefore facilitates easy access and increased consumption’’* (a resident of the village of Kampouya). An additional reason for the increase in *M. oleifera* consumption is the reduction in accessible alternative food sources. There are very few vegetables accessible in the dry season apart from *M. oleifera* and *okra* (*Abelmoschus esculentus)*. In addition, we noticed temporal aspects that enhanced *M. oleifera* consumption since storage and processing extended the consumption period to the whole year.

Regarding the **affordability and convenience dimension**, sufficient quantities of the leaves were easily found in the natural environment, sometimes from the free transfers that made them affordable and convenient. On the other hand, the small size of the leaves made them easy to collect and cook.

For the **desirability dimension**, a **better awareness of the usefulness**, mainly about health and nutrition, as well as new processing, storage, and preparation techniques have been declared. Although this species has long existed in the research area, information on the edibility and usefulness of its leaves was unknown during most of the participants’ youth. Fresh or dried leaves were used to treat various illnesses and problems (malaria, tuberculosis, AIDS, stomach pains, breastfeeding problems in women, malnutrition). Some interviewees were aware of the edibility of the leaves but consumed them to a lesser extent than today: *Ten years ago, no, I’d say 15 years ago, I had Moringa in my neighborhood and even in my house. But we didn’t eat the leaves. So, people from Niger and Mali came to my house, always asking for the leaves. But today, I eat a lot of it and even have some at home. It’s become so much more popular than before, as people have discovered its many uses and virtues* (a resident of the village of Kampouya). Then, leaves were partially used as animal fodder during the interviewees’ youth. Information on the nutritional and health benefits of *M. oleifera* leaves has been reinforced by the promotion activities of various stakeholders, including local projects, hospitals, scientists, start-ups, and the media. These activities have raised local awareness of the leaves’ dietary benefits by recommending them as part of a recovery diet for malnourished children, by marketing products made from the species’ leaves or through information campaigns; the storage of the plant’s leaves in powder form has helped to increase consumption

Another aspect of desirability, such as **preferences for sensory properties** contributed to the positive evolution of consumption. For example, *M. oleifera* became increasingly recognized and known as an edible species, and the formerly wild species was increasingly cultivated. Furthermore, the **social marker** aspect had a positive influence on the upward trend of *M. oleifera* consumption since their image changed from a foreign element to a more familiar and appreciated element of the human diet.

#### 3.4.2. Adansonia digitata

A fluctuation trend can be observed for *A. digitata* parts (a decreasing trend for the pulp and a varying trend for the kernels and leaves). For the participants in this study, the decreasing trend of *A. digitata* pulp could be explained by several dimensions of the personal food environment. Their **low accessibility** was due to insufficient reproduction efforts by humans combined with a decreased productivity of elder trees due to the long lead time required to start harvesting fruit (*A. digitata*). This perennial species was destroyed by cutting or bushfires. In addition, the increased production of a variety of alternative food sources has reduced consumption of *A. digitata* pulps. Indeed, the population depended less on the wild harvest of these fruits as they substituted them with the production of other plant species such as *Dioscorea L.* (yams), *Oryza sativa L.* (rice), and *Glycine max (L.)* (soja).

The **affordability dimension** in terms of purchasing power partially impeded or reduced the purchase and consumption of *A. digitata* pulp. Indeed, baobab pulp is one of the edible plants with a high economic value. It is therefore marketed, and the cash income from sales is used to buy food (e.g. staples, condiments) and other items (e.g. soap, washing powder, shoes, clothes). As for the leaves and kernels, for example, the increasing sales have harmed auto consumption by the population.

In terms of **convenience dimension**, the acquisition of the pulp is a factor impeding consumption due to the unpractical harvesting, essentially linked to the high height of the baobab trees: ‘‘*Can you climb the baobab that easily? No, you can’t, unless it’s a young baobab. Then you’ll have to wait for the rainy season to start, with the high winds, before you can expect the fruit to fall’’* (a resident of *Tectibayaou*’s village). Although people were aware of how to **process and store** baobab leaves throughout the year, there were still problems with hygiene standards and preserving the nutritional value of the leaves. For example, clean drying areas were often insufficient, yellow leaves were often not removed, and drying often proceeded in the sun. There is also a lack of material inputs (e.g. refrigerator, clean water, bottles) to process and store *A. digitata* pulp.

Regarding the **desirability dimension**, **informal institutions** in terms of decreased **social access** regarding harvest and transfers contributed to the downward trend of *A. digitata*’s pulps. In the past, *A. digitata* fruit that fell to the ground could be harvested by anyone, but it is becoming increasingly common for the tree’s owner to restrict access. In addition, a loss of familiarity and changes in dietary habits were contributing to the decline in consumption. For instance, mothers and grandmothers were used to prepare porridges with the pulp which nowadays stopped. Certain **taboos** in the *Berba* and *Waama* ethnic groups impeded the consumption of baobab. According to them, anyone who planted an *A. digitata* tree and consumed its fruit, leaves, or kernels was condemned to die. The reproduction of the trees was perceived as being reserved for God, even if people were allowed to maintain existing trees: ‘‘*You know, some baobabs are possessed by spirits, souls. Some old people can even feel these spirits. In such cases, they forbid their consumption at the risk of death’’* (a resident of Natitingou’s village). The absence or limited reproduction of perennial species as well as the **lack of knowledge** of their reproduction methods and the mistaken belief that these species only reproduce naturally could lead to a reduction in consumption. Furthermore, a decrease in **sensory preferences** related to unfavorable taste perceptions such as sourness of *A. digitata*’s pulp or bitterness of old leaves.

#### 3.4.3. Vigna radiata

Essentially, it was **dimensions of convenience** and **desirability** that had positive influences on the upward trend of *V. radiata* seeds. A few years ago, with the arrival of the variety, convenience became greater due to its shorter production cycle and shorter cooking time: ‘‘*If the first rains fall and you plant the species, it will produce about two months later. If you wish, you can plant a second seedling during the same season’’* (a resident of Kosso’s village); ‘‘I’ve *been producing the species for 3 years, it’s only this year that I haven’t been able to because of my pregnancy. I love its smell and taste’’* (a resident of Douani’s village). Most participants who recognized this species said it tasted very good and was much appreciated. Moreover, various known techniques for preparing *V. radiata* seeds, which were similar to other legumes such as *V. subterranea* (bambara peanut) or *V. uniguculata* (cowpea), increased their consumption ‘‘In *our cooking demonstrations, we used mung bean instead of cowpea to enhance meals. We use it to make fritters, cakes and even Watché (Rice + Beans)*’’ (an agent working for Giz/Prosar).

#### 3.4.4. Ocimum gratissimum

A predominant trend of consumption of *O. gratissimum*’s leaves was marked by variation of the consumption. In fact, for **physical accessibility**, herbs like *O. gratissimum* were accessible through their wild and cultivated growth around houses. Its seeds are purposively thrown away in the fields after the harvest of their leaves to enhance their natural reproduction. *O. gratissimum* mainly grew in humid locations. Watering of women with kitchen water at drier locations increased its production and consumption. The increasing consumption was also due to its **affordable** option compared to some vegetables which can be acquired expensive. Its preparation was found to be highly convenient due to the short amount of time needed. Positive **social maker** (**high appreciation)** of the leaves referred to their importance in dietary habits/as part of the local culture.

Then, **awareness of the health benefits** of the consumption of plants part showed positive effects on the consumption. Indeed, the leaves have antibiotic effects mainly used to treat stomach-related diseases (e.g. intestinal infections, constipation, diarrhea, or ulcers). It constituted the first sauce that was given to women after giving birth because of its stimulating effect on milk production ‘‘ *Yes, Ocimum can be eaten, but it’s better known as a therapeutic food* ‘‘(a resident of Koukorbirgou’s village). In addition, the smell had the positive effect of reducing the undesired odor of other food, although, for new consumers of the species, the strong smell represented a barrier to consumption. Several obstacles have also been mentioned, leading to fluctuations in consumption over time. Indeed, the limited reproduction was explained by **a lack of knowledge** about their reproduction modes, by a lack of interest or incentives**. Lack of storage** and **processing techniques** or the willingness to find them decreased the leaves consumption.

## 4. Discussion

Regarding the investigated period, the results showed that not all species became less consumed by the research sample. Their consumption was not declining among all plant parts but showed upward, downward, and varying trends. The changes in consumption patterns of the investigated NUS were influenced by various factors across all aspects of the food environment. While sociocultural factors were found to drive declines in consumption in some studies predominantly (22), others identified accessibility and environmental factors as key drivers (23).

In this study, it has been found out an uptrend for *M. oleifera* leaves and *V. radiata* seeds consumption. *M. oleifera* is a traditional leaf vegetable that is widely distributed geographically in Benin (24). This wide distribution has probably resulted from its adaptation to the different ecological conditions and its wide climatic tolerance. Indeed, *M. oleifera* was only introduced in Africa at the beginning of the 20^th^ century and was therefore not yet anchored in family and cultural practices (25). Several people interviewed said they were not aware of the uses of moringa in their youth, unlike today. Other studies in Benin show that the local naming of *M. oleifera* in rural communities in Southern Benin can be interpreted as its use for food and medicinal purposes is relatively recent (26). It is an alicament. Then, the interviewees explained the positive effect of accessibility by the fact that it is growing more and more in home gardens. Agoyi *et al.* (2014) reported that the most common *M. oleifera* growing systems encountered in southern Benin were individuals scattered around farms, home gardens, and fences. It is therefore an easy species to find and can be harvested easily and free of charge in the wild or by free transfer. About **temporal aspects**, like *okra*, *M. oleifera* is also one of the food species accessible in seasons of food scarcity. The same was founded in Ghana, where *M. oleifera* leaves and several other green leafy vegetables as African spinach (*Amaranthus hybridus)*, amaranth (*Amaranthus cruentus L.)*, leafy eggplant (*Solanum melongena L.*), and jute mallow (*Corchorus olitorius*) are available and accessible all year round (27). An interesting finding in this study is the consistent impact of food and nutritional knowledge and skills uniformly contributing to the upward trend of *M. oleifera* leaves. This proves that just as in South Africa (28), several sources such as the media, schools, and research have communicated the importance of this plant in the diet of populations in the study area. Raising awareness of Moringa’s multiple uses throughout the community is necessary to create market demand and maximize resource utilization (29).

Whereas some studies identified taste as the most important sociocultural reason for the increase in wild edible plants’ consumption (22), positive influences of taste preferences were mentioned mostly for *V. radiata’s* seeds. A study conducted by Bankole *et al.* (2023) showed that 83.33% and 82.69% of children respectively appreciated mung bean boiled with moringa leaves (MBML) and mung bean boiled with roasted maize flour (MBMF). Anggraeni & Rahmawaty (2022) in they study also found that the higher the addition of mung bean flour, the more panelists preferred brownies suweg formulated. As consumption has significantly increased over time, participants have observed that certain convenience-related factors have made the harvesting of this species challenging. Indeed, itchy hairs during harvest and the small size of the seeds make harvesting difficult. A study conducted by Sequeros *et al.* (2021) emphasized the improvement of mechanization to maintain profitability as rising labor costs depress *V. radiata* production. Market accessibility is limited due to low levels of production. In Kenya, for example, unstable markets and low market prices have been observed due to a lack of clear information on *V. radiata* prices set by traders, which do not always reflect the reality for producers (32).

Many of the causes of the decrease in accessibility of *A. digitata* pulp were similar to the findings of another study in Benin which explained that it was due to limited regeneration caused by biotic and abiotic disturbances (33). Then, several studies indicate that socio-economic reasons contribute to the decline of NUS (Hunter et al., 2019). It is clear that wild plants continue to represent an important part of the global food basket, while a variety of social and ecological factors are acting to reduce the use of wild foods, their importance may be set to grow as pressures on agricultural productivity increase (34). However, increased access to markets may also reduce their consumption (35). The current decrease in consumption levels of *A. digitata* pulps is a result of their increasing sale. Other studies further support the high sales levels of the plant parts (36). In contrast to the pulps, it is interesting to notice that the leaves consumption has not been negatively impacted by their marketing. It is worth mentioning that the leaves are among the ten most traded traditional vegetables in the Sudanian zone of Benin, and nationally as well (37). Additionally, they are also exported to other West African countries (38). *A. digitata* pulp has also suffered from the difficulty of processing and storage. The study by Chadare *et al.* (2008) showed that in Benin, most ethnic groups mentioned the difficulty of certain processing operations, in particular seed hulling, grinding, and sieving operations, as well as the problems of storing and preserving almonds and pulp.

Aspects such as negative beliefs and image, taboos were seen as barriers to NUS consumption (23,39). According to the interviewees, the taboo on regenerating *A. digitata* had religious and cultural reasons. Other studies confirm the important socio-cultural role of *A. digitata* in local belief systems and for cultural identity in Benin (38,40). Reducing free transfers in the close social environment affected the consumption of different parts of *A. digitata* which suffered from negative side-effects of their high sales level.

As for *O. gratissimum*’s leaves, inconvenient acquisition contributes to decreased consumption of the *O. gratissimum leaves.* Indeed, **lack of processing and storage facilities** as well as a **lack of knowledge** about their reproduction modes contributed to the reduction of leaves consumption. Dossoukpevi *et al.* (2012) pointed out that climate change is hampering the good productivity of the leaves through traditional production methods (seed sowing, vegetative propagation), affecting their availability and consumption. Baco’s (2019) study revealed that some households mentioned a lack of knowledge, organoleptic attributes, and totemic considerations to justify the non-consumption of certain leafy vegetables, including African basil (*O.gratissimum*). Similar reasons were noted in several other studies for the non-consumption of certain traditional leafy vegetables (43). In other respects, an interesting find in this study is the consistent impact of food and nutritional knowledge and skills were uniformly contributing to the varied trend as well. A study confirms the strong link of *O. gratissimum* consumption to medicinal use awareness in terms of antibiotic properties (37). Dansi *et al.* (2008) study also showed that consumption was higher during menstruation, pregnancy, and breastfeeding. The therapeutic properties were multiple and linked to the leaves’ ability to eliminate blood clots, facilitate childbirth, eliminate waste after delivery, treat post-partum infections, heal wounds, and stimulate milk secretion. Reducing free transfers in the close social environment affected consumption of *O. gratissimum* which suffered from negative side-effects of their high sales level.

### 3.2. Limits of the study and recommendations

A large number of barriers and enhancers have been identified in this work, but not all of them likely determine consumption to the same extent. Overall, the intensity of influences and their interrelationships, including trade-offs between dimensions of the personal food environment have not been fully considered. Theoretical and methodological refinements are needed to measure the other different types (natural/built), domains (internal/external), and parameters (dimensions and aspects) of food environments (16) for a better understanding of the factors influencing NUS consumption.

In terms of recommendations, we strongly suggest involving local communities in the promotion of neglected and underutilized species (NUS) to ensure their conservation and sustainable use. In food systems where producers and consumers intersect, involving smallholders is crucial for preserving plant species (44). Protecting on-farm and wild species and their ecosystems is vital, given the reliance on natural environments for NUS acquisition. Enhancing access to market presence can improve the availability of specific plant parts. Educating about the nutritional benefits of these plants is essential (45). Strengthening local social networks acknowledges their dynamic nature and fosters community cohesion. Empowering women with leadership roles and decision-making authority promotes inclusivity (46). Strategies should address taboos and misconceptions related to plant consumption, aiming to foster positive perceptions and increase consumption.

## Conclusion

This study showed that among all species investigated, *A. digitata* and *M. oleifera* were the most consumed and preferred species. Consumption is not automatically declining for all species. Indeed, increased trends in consumption were found for *M. oleifera* leaves and *V. radiata* seeds while *A. digitata* pulp consumption decreased. The results provide useful insights for developing effective entry points for improved consumption of the NUS studied, by characterizing barriers and improvement factors in the various sources of acquisition and consumption processes. Some plant parts have barriers to overcome in all four dimensions of the local food environment, whereas only some dimensions are challenging for others. More research and supportive measures in all four dimensions of the personal food environment are needed to create supportive food environments.

## Acknowledgments

The authors wish to thank GIZ and the CGIAR FRESH initiative for their support. Special thanks go to the residents of the study villages, who facilitated the implementation of the activities.

